# Whole Genome Sequences of *Aedes aegypti* (Linn.) Field Isolates from Southern India

**DOI:** 10.1101/2020.05.08.083949

**Authors:** Venitha Bernard, Sanjana Moudgalya, Daniel Reegan, Peddagangannagari Sreekanthreddy, Abhilash Mohan, Hosahalli S Subramanya, Shruthi Sridhar Vembar, Sanjay Ghosh

## Abstract

Aedes spp. mosquitoes are a major health concern as they transmit several viral pathogens resulting in millions of deaths annually around the world. This is compounded by the emergence of insecticide-resistant strains and global warming, which could expose more than half of the world's population to Aedes-borne diseases in the future. Therefore, a comprehensive understanding of vector biology and the genomic basis of phenotypes such as insecticide resistance in natural populations are of paramount importance. Here, we sequenced the genome of Aedes aegypti mosquitos sampled from dengue-endemic areas and investigated the genetic variations between the previously reported laboratory-reared strain and our field isolates. The mosquito genomic DNA was used for paired-end sequencing using the Illumina platform. The reads were used for template-based assembly and mapped to the Aedes aegypti reference genome. Stringent parameters and multiple variant calling methods were used to identify unique single nucleotide variants (SNVs) and insertions-deletions (indels) and mapped to the Aedes chromosomes to create a draft consensus genome. Gene Ontology analyses was performed on the variant-enriched genes while two gene families involved in insecticide resistance were used for comparative sequence and phylogenetic analyses. Comparative sequence variant analyses showed that the majority of the high-quality variants in our samples mapped to non-coding regions of the genome, while gene ontology analyses of genic variants revealed enrichment of terms relevant to drug binding and insecticide resistance. Importantly, one mutation implicated in pyrethroid resistance was found in one Aedes sample. This is the first report of genome sequences of A. aegypti field isolates from India which reveals variants specific to the wild population. This is a useful resource which will facilitate development of robust integrated vector control strategies for management of Aedes-borne diseases through genetic manipulation of local mosquito populations.

## INTRODUCTION

*Aedes spp*. are the principal mosquito vectors for transmission of etiological agents of arboviral diseases such as yellow fever, chikungunya, dengue, and zika [1]. Together, these viral diseases globally infect up to 500 million people every year with increasing fatalities and significantly contribute to the public health burden of several countries [2,3]. *Aedes* mosquitoes are found in tropical and sub-tropical climates across all continents, while predictions show that climate change will increase the habitat and distribution of *Aedes* which could expose a larger human population to the risk of *Aedes*-borne diseases [4,5,6].

In India, *Aedes aegypti* and *Aedes albopictus* are the major vectors for Dengue and Chikungunya, especially around the monsoon season [7]. Dengue, in particular, is endemic to all states in the country with high occurrence in urban, peri-urban and rural areas [8] often resulting in significant morbidity and fatality. In the absence of effective vaccines and therapeutic treatments against arboviruses, the current control strategy for these diseases relies heavily on targeting *Aedes spp*. populations to regulate disease transmission. However, traditional mosquito suppression methods using pesticides is ineffective and has become unsustainable as mosquitoes develop resistance quickly [9]. Therefore, there is a global demand for developing alternative safe and effective approaches to target vector competency based on a molecular understanding of the insect vector.

Whole genome sequencing has emerged as a powerful method to reveal genetic variations as well as provide crucial insights into the biology of several organisms including *Aedes* [10,11]. While the first *A. aegypti* genomes were reported in 2007 [12], a high quality, chromosome-level reference genome of an *A. aegypti* laboratory strain LVP_AGWG (referred to as AaegL5) with a genome size 1.28 Gb was reported recently by the Liverpool Aedes Genome Working Group [11]. The authors used Single Molecule Real-Time sequencing of long reads and chromosome conformation capture to map a majority (>94%) of the sequenced reads to three *Aedes* chromosomes leading to fewer gaps in the primary assembly. The improved genome not only revealed new genes and gene families involved in human host seeking and insecticide receptors but also provided insights into the genetic basis of resistance to pesticides [11]. Furthermore, a recent comparative study using whole genome sequence data of field isolates of *A. aegypti* in California demonstrated significant genetic differences between geographical isolates and revealed genomic signatures of potential selection, thus providing insights into population dynamics and mosquito evolution [13]. These studies highlighted the importance of genome sequence data in providing insights into the vector biology. However, genome sequence information of *Aedes* field isolates from India is currently unavailable.

*A. aegypti* is the dominant species in the city of Bengaluru in south India which is endemic to dengue infections [14]. To understand the genetic diversity of the natural *A. aegypti* populations and gain an insight into vector of competence in India, we sequenced the genomic DNA of two *A. aegypti* female mosquitoes (called AEBAN1 and AEBAN2) collected from different locations in Bengaluru and analysed their genomes in reference to the AaegL5 assembly. Our analysis revealed the existence of several high-quality variants in AEBAN1 and AEBAN2 that could contribute to divergent genetic architecture of *A. aegypti* natural populations in India. To our knowledge, this is the first report of genome sequences of *Aedes* field isolates from India. This is a useful resource for the research community to perform comparative genomic analyses and will aid future studies on vector competence of mosquito populations from arboviral-endemic areas.

## METHODS

### Sample collection, preparation of genomic DNA library and Illumina sequencing

The larvae of *Aedes* mosquitoes were collected from two localities in Bengaluru city, namely Koramangala (12.9352° N, 77.6244° E; called AEBAN1) and Konappana Agrahara (12.9679° N, 77.5625° E; called AEBAN2). The larvae were transported to the laboratory within 2 h and fed with an artificial feed made of dog biscuits and yeast (60:40 ratio). The adult mosquitoes that emerged were identified as *Aedes aegypti* (Linn.) and the females were used for further analyses. For each location, genomic DNA (gDNA) of a single adult female mosquito was isolated using the DNeasy Blood and Tissue kit (Qiagen # 69504) following manufacturer instructions. ~200 ng of gDNA was used for library preparation with the NEBNext DNA Ultra II Library Prep Kit (New England Biolabs # E7645S) as per manufacturer’s instructions. The final library was quantified by Qubit 2.0 fluorometer (Thermo Fisher Scientific), further verified by qPCR using NEBNext Library Quant Kit for Illumina (New England Biolabs #E7630S) on Stratagene Mx3000P instrument and its profile analysed by Agilent Tape station 2200. Mean fragment size of the library for AEBAN1 and AEBAN2 were 450 bp and 395 bp, respectively. Denatured libraries were clustered on cBot (Illumina) and analyzed by paired-end sequencing using the Illumina HiSeq 2500 with v3 chemistry for 209 cycles: the read configuration was 101+7+101. Base call files were demultiplexed to FASTQ format using bcl2fastq v2.18.0.12. Greater than 80% of the bases had phred quality scores above Q30.

### Quality control, mapping and preprocessing of DNA sequences

The FASTQ files of both samples were checked for quality using FASTQC v0.11.5 [15]. Sample AEBAN2 was trimmed using Trim Galore! [16] v0.5.0 in paired-end mode, with other parameters set at default. The raw FASTQ files of AEBAN1 and the trimmed FASTQ files of AEBAN2 were then mapped to the AaegL5 assembly of the *A. aegypti* reference genome for strain LVP_AGWG [11], which was downloaded from VectorBase [17]. The mapping was performed using BWA-mem v0.7.12-r1039 [18]. The output SAM files were converted to BAM format and sorted by coordinate using Picard’s SortSam v2.18.26-SNAPSHOT (http://broadinstitute.github.io/picard). BAM statistics were then extracted using the Java-based QualiMap v2.2.1 [19]. Duplicates in the BAM file were marked using Picard MarkDuplicates module. BAM index files (.bai) and dictionary files (.dict) were also created using Picard, through BuildBamIndex and CreateSequenceDictionary modules, respectively. Reference genome index (.fai) was created using samtools faidx, v0.1.20 [20]. Mapping statistics of the samples are provided in **Table 1**.

**Table 1:** Mapping statistics of the fastq files generated for AEBAN1 and AEBAN2

| Mapping Statistic | AEBAN1 | AEBAN2 |
| --- | --- | --- |
| Number of reads | 433,751,605 | 97,807,809 |
| Mapped reads | 414,159,840 /<br>95.48% | 95,479,563 /<br>97.62% |
| Unmapped reads | 19,591,765 /<br>4.52% | 2,328,246 /<br>2.38% |
| Mapped reads, first in pair | 207,471,939 /<br>47.83% | 47,760,767 /<br>48.83% |
| Mapped reads, second in pair | 206,687,901 /<br>47.65% | 47,718,796 /<br>48.79% |
| Mapped reads, both in pair | 407,704,571 /<br>93.99% | 94,550,716 /<br>96.67% |
| Mean Coverage | 30.4126 | 6.9237 |
| Mean mapping quality | 12.44 | 12.4 |
| Median insert size | 284 | 199 |
| GC% | 38.73% | 38.34% |
| Number/percentage of N's | 7,473,793 /<br>0.02% | 161,163 / 0% |

### Variant calling

Single Nucleotide Variants (SNVs) and Insertions-Deletions (Indels) in the AEBAN1 and AEBAN2 sequences were identified using three different variant callers, GATK v3.8-1-0-gf15c1c3ef [21], FreeBayes v1.2.0-17-ga78ffc0 [22], and Platypus [23]. In GATK, after indel realignment, a first round of variant calling was performed using the module HaplotypeCaller. These were then processed by the BaseRecalibrator module, after which a second round of variant calling was performed. This resulted in a final set of SNVs and indels, which were filtered and merged into a single VCF file for further analysis. In FreeBayes, variants were called using the sorted and deduplicated BAM files. For AEBAN1, variants with QUAL > 20 and sequencing depth (DP) > 20 were retained while for AEBAN2, variants with QUAL > 60.2 and DP > 7 were retained. In Platypus, variants were called using default parameters. Additional filtering was performed similar to FreeBayes.

### Consensus genomic sequence creation

To identify high confidence variants in each sample, an intersection of the results from GATK, FreeBayes and Platypus was generated using bcftools v1.9-158-gf24ccc9 (https://github.com/samtools/bcftools) (**Fig. 1A**). This yielded 3157499 & 1069359 SNVs for AEBAN1 and AEBAN2 respectively, and no indels. Therefore, for indels, the intersection between GATK and Platypus was considered, which yielded 673730 and 150128 indels for AEBAN1 and AEBAN2, respectively (**Fig. 1A**). These were merged with the common SNVs into a final VCF file that contained biallelic and multiallelic variants; for the latter, only those alleles that had the highest allelic depth (AD) were retained. This VCF file was then used to create the final consensus genome for AEBAN1 or AEBAN2 using an updated version of bcftools (v1.9-221-gb1426c5) and the *A. aegypti* AaegL5 genome [11] as template. The consensus genomes have been submitted to the NCBI Genome database. Variant density plots across the *A. aegypti* chromosomes for AEBAN1 and AEBAN2 were generated using bedtools version v2.29.2 [24] and plotLy (https://plotly.com/).

**Figure 1.**
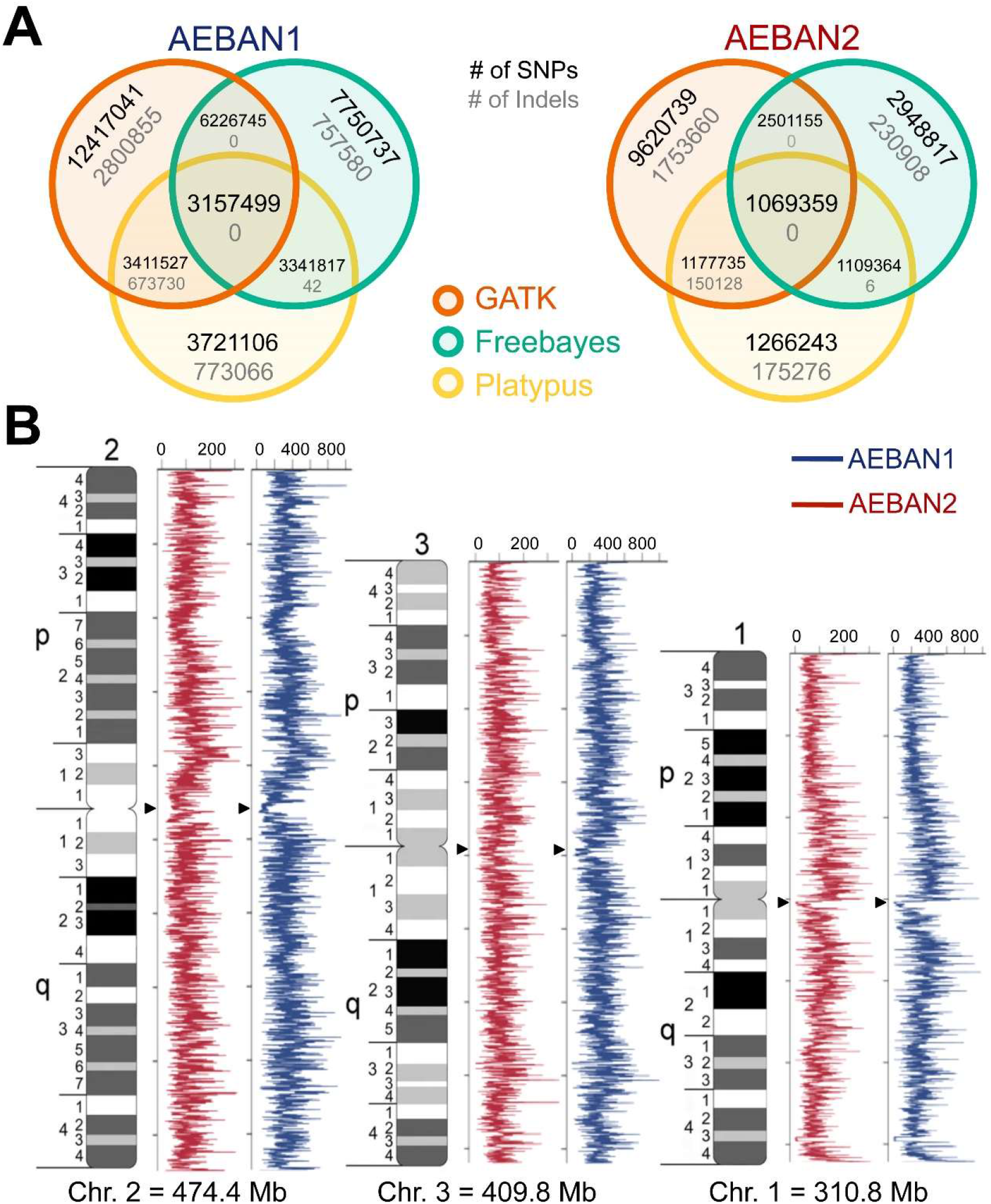
The genomes of AEBAN1 and AEBAN2 are divergent from AaegL5. *(A)* Venn diagrams showing the intersection of SNVs (black) and Indels (gray) in AEBAN1 and AEBAN2 relative to AaegL5 using three variant callers GATK (orange), FreeBayes (green) and Platypus (yellow). *(B)* Cytogenetic maps showing the variant density of AEBAN1 (blue) and AEBAN2 (red) along the three *A. aegypti* chromosomes. The variant density across 100 kb bins of the three chromosomes was calculated using bedtools v2.29.2 [24] and plotted for the AEBAN1 and AEBAN2 samples (Adapted from [32]). Arrowheads indicate the position of the centromere for each chromosome.

### Nucleotide sequence accession numbers

The genome sequences of the *A. aegypti* isolates analysed in this study have been deposited at DDBJ/EMBL/GenBank under the accession numbers WELD00000000 (AEBAN1) and WELE00000000 (AEBAN2).

### Gene ontology analysis

For Gene Ontology (GO) analysis, variants localizing to the three *A. aegypti* chromosomes were extracted from the final VCF files and annotated using SnpEff (v4.3t,2017-11-24) [25]. A customized database was built for annotation using the AaegL5.2 gene set [17]. GO enrichment analysis was performed using GOEnrichment [26] through the Galaxy v2.0.1 server at usegalaxy.org [27]. AaegL5.2 GO mappings were downloaded from VectorBase BioMart (https://www.vectorbase.org/resources/biomart), and core GO terms from the GO Consortium website (http://purl.obolibrary.org/obo/go.obo). Multiple test correction of enriched GO terms was performed using the Benjamini-Hochberg method [28] with a p-value of 0.05. Other options were left at default.

### Phylogenetic analysis of VGSC and Cys-loop LGICs

For phylogenetic analysis, indel-free AEBAN1 and AEBAN2 genomes were reconstructed by using only the SNVs called using the three different methods, *i.e.*, 3157499 SNVs for AEBAN1 and 1069359 SNVs for AEBAN2. Next, the coding sequence (CDS) of the VGSC gene (NCBI Nucleotide ID XM_021852342.1) from the AaegL5 assembly was used to extract the orthologous CDSs from AEBAN1 and AEBAN2 using GMAP v2020-03-12 [29]. Lastly, the VGSC protein sequence of AaegL5 (NCBI Protein ID XP_021708034) along with the predicted protein sequences of VGSC from AEBAN1 and AEBAN2 were aligned using MUSCLE in MEGA X [30]. The multiple sequence alignment (MSA) was visualized using MView [31].

The AaegL5 CDSs of fourteen nicotinic acetylcholine receptors (nAChr) α1-10 and β1-4, belonging to the Cys-loop Ligand-Gated Ion Channel (LGIC) family were used to extract orthologous protein sequences from the indel-free AEBAN1 and AEBAN2 genomes. MSA of these sequences, along with the Erwinia ligand-gated ion channel (ELIC) from *Dickeya chrysanthemi* (NCBI Protein ID P0C7B7) as the outgroup, was performed using MUSCLE. The best protein substitution model for the data was selected by maximum likelihood in MEGA X using default parameters and was found to be the Le and Gascuel model (LG) with gamma distributed sites (G) (data not shown), The phylogenetic tree was constructed by using the above model and 500 bootstrap replications. The gaps/missing data were treated as partial deletions with the site coverage cut-off at 95%. All other parameters were left at default.

## RESULTS

For genomic analyses, we extracted genomic DNA of *A. aegypti* mosquitoes collected from two independent locations in Bengaluru (herewith referred to as AEBAN1 and AEBAN2) and processed the DNA for Illumina sequencing-by-synthesis. The resulting high-quality sequencing data (Phred quality score >30) were used for template-based genome assembly with the *A. aegypti* AaegL5 reference genome [11] as template. The sequencing reads were mapped to the reference genome by aligning the fastq files of AEBAN1 and AEBAN2 to AaegL5 assembly (**Table 1**). Despite differences in the coverage, the vast majority of the sequencing reads of both samples mapped to the AaegL5 genome (AEBAN1 coverage 30x, 95.48% mapped reads; AEBAN2 coverage 7x, 97.62% mapped reads).

To identify the genomic differences (SNVs and indels) between the laboratory strain AaegL5 and field isolates AEBAN1 and AEBAN2, we used three different variant callers: Genome Analysis Toolkit (GATK), Freebayes, and Platypus, to obtain high-quality genomic variants (see Materials and Methods for details). The numbers of SNVs and indels identified by each method are shown in **Fig. 1A**. Only SNVs common to all three methods (31,57,499 for AEBAN1 and 10,69,359 for AEBAN2), and indels identified by both GATK and Platypus (6,73,730 for AEBAN1 and 1,50,128 for AEBAN2) were used to assemble the final consensus genomes. Notably, the draft genomes of AEBAN1 and AEBAN2 are 1278422764 bp and 1278641819 bp, respectively. Compared to the *A. aegypti* AaegL5 reference genome, these genomes are smaller by 3,09,340 and 90,285 bp, respectively, primarily due to the structural variations such as of deletions.

We next analyzed the distribution of variants across the three *A. aegypti* chromosomes for both samples. As shown in **Fig. 1B**, the variants did not cluster at any particular region of the chromosomes, rather they were distributed quite evenly across the p and q arms of each chromosome. However, we observed elevated variant density at select regions within each chromosome suggesting that these DNA segments in AEBAN1 and AEBAN2 might be more prone to recombination and/or accumulation of mutations. The lowest variant density was found around the centromeres (**Fig. 1B**; arrowheads) - such a decline in genetic diversity at centromeres has previously been observed for *A. aegypti* [11] and implies reduced recombination and/or mutation rates in this region. Importantly, the overall pattern of variant distribution for AEBAN2 was comparable to that of AEBAN1 indicating that the lower sequencing depth of AEBAN2 did not compromise our ability to identify genomic variations relative to the *A. aegypti* AaegL5 reference.

To determine the distribution of variants across different genomic regions of AEBAN1 and AEBAN2, we grouped the SNVs and indels that mapped to the three *A. aegypti* chromosomes into 5’UTR, Exonic, Intronic, 3’UTR or Intergenic categories using SnpEff [25]. As shown in **Fig. 2A** and **Supplementary Table S1**, in both samples a vast majority of the variants mapped to the introns (73.95% in AEBAN1, 73.41% in AEBAN2) and intergenic regions (11.97% in AEBAN1, 11.50% in AEBAN2) of the genome suggesting minor perturbations in the proteomes of AEBAN1 and AEBAN2 relative to AaegL5. Given the non-coding regions of the genome contain structural, regulatory and transcribed information, variants mapping to these regions of the genome may contribute to altered regulation and/or expression of protein-coding genes, thus impacting phenotypes including insecticide resistance and vector competence.

**Figure 2.**
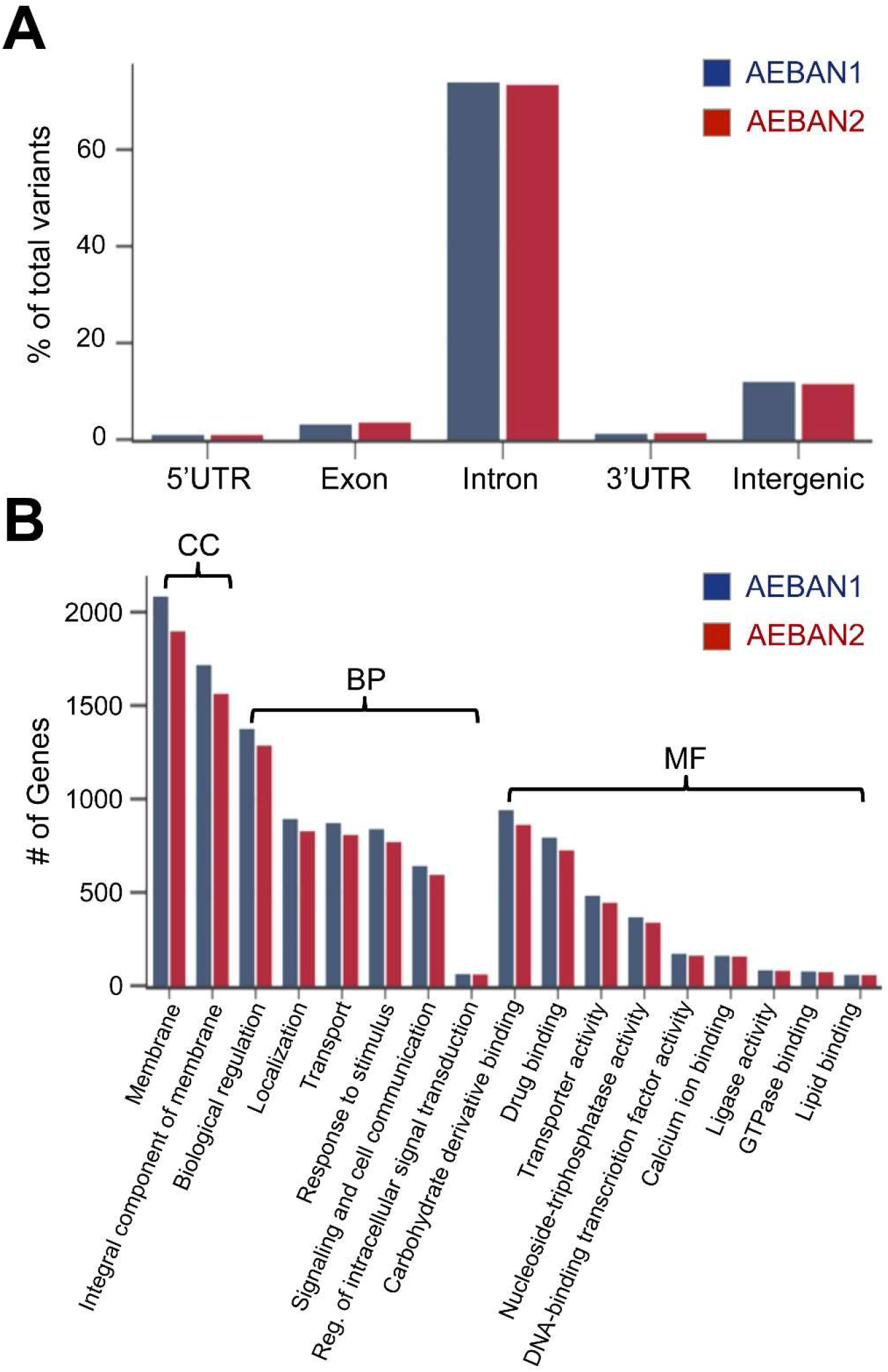
Features of variants identified in the AEBAN1 and AEBAN2 genomes. *(A)* The percentage of variants mapping to different genomic regions of the three *A. aegypti* chromosomes for AEBAN1 (blue) and AEBAN2 (red) are shown. (See Table 1 for details). *(B)* Gene ontology (GO) analysis of the variants reveals significantly enriched GO terms. The numbers of genes associated with each term are shown for the Cellular Component (CC), Biological Process (BP) and Molecular Function (MF) categories for AEBAN1 (blue) and AEBAN2 (red).

To assess the impact of genomic variants on mosquito biology, we performed Gene Ontology (GO) analysis using all variants that mapped to the 5’UTR, exonic and 3’UTR regions of the chromosomes. This analysis revealed several GO terms that were enriched (False Discovery Rate <= 0.05) and common to both AEBAN1 and AEBAN2 (**Supplementary Table S2**). The most significant amongst these are as follows: (a) in the Cellular Component (CC) category - membrane and integral component of membrane, (b) in the Biological Process (BP) category - biological regulation, localization, transport, response to stimulus, signaling and cell communication, and regulation of intracellular signal transduction, and (c) in the Molecular Function (MF) category - carbohydrate derivative binding, drug binding, transporter activity, nucleoside-triphosphatase activity, DNA-binding transcription factor activity, calcium ion binding, ligase activity, GTPase binding and lipid binding (**Fig. 2B**). Importantly, enrichment of GO terms like drug binding and transporter activity indicates that the field isolates of *A. aegypti* mosquitoes from Bengaluru might have adapted to insecticides used widely for vector control programs locally and in India.

To explore further, we analyzed SNVs in two gene families - the voltage-gated sodium channel (VGSC), a known target of pyrethroid insecticides [11,33] and fourteen nicotinic acetylcholine receptors (nAChr) α1-10 and β1-4 of the Cys-loop ligand-gated ion channel (Cys-loop LGIC) family, which are being developed as larvicidal targets [11,34]. While the 5 SNVs in the sequence of AEBAN2 VGSC did not alter amino acid sequence as compared with AaegL5 VGSC, the coding sequence of VGSC from AEBAN1 showed 16 SNVs resulting in three amino acid changes, namely S989P, V1016G and V1704G (**Supplementary Fig. S1**). Importantly, mutation of valine at position 1016 to isoleucine (V1016I) has previously been reported in natural *A. aegypti* populations that are resistant to pyrethroids [35,36,37]. This observation suggests that the Indian *aegypti* populations might be developing resistance to pyrethroids. Notably, phylogenetic analysis of nAChr protein sequences from AaegL5, AEBAN1 and AEBAN2 revealed similarities to as well as divergence from AaegL5 (**Fig. 3)**. For instance, the nAChr-α3, α4, α7 and β3 proteins of AEBAN1 and AEBAN2 are more similar to each other as compared to their AaegL5 counterparts while the nAChr-α2, α5, α6 and β1 proteins of AEBAN1 and AaeGL5 are more similar to each other as compared to their AEBAN2 counterparts. Taken together, these analyses highlight the genomic diversity between the AaegL5 reference strain and field isolates from Bengaluru and emphasize the need to use population genomics data to design optimal vector control strategies in a country/region-specific manner.

**Figure 3.**
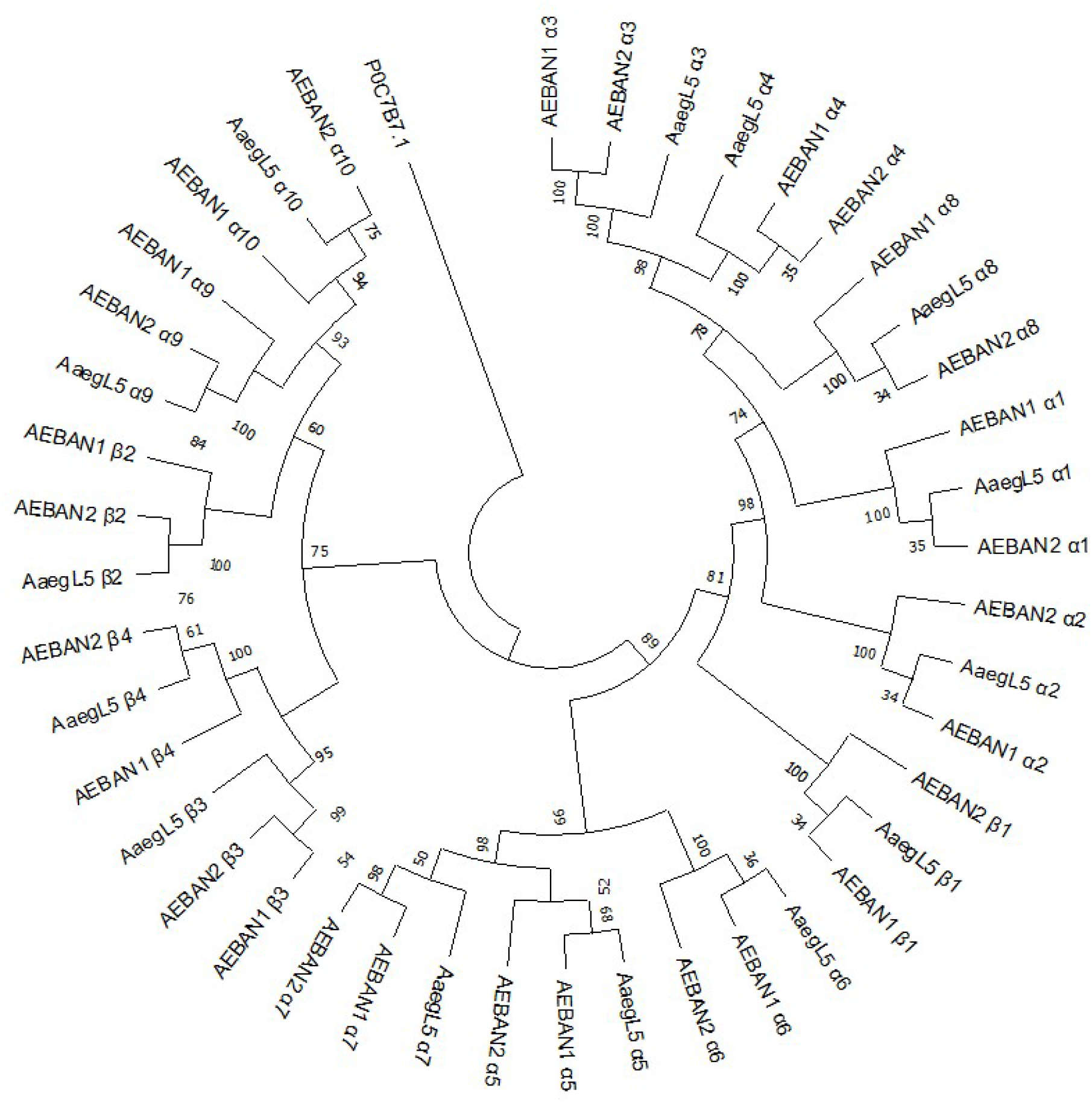
Phylogenetic analysis of nAChr α1-10 and β1-4 proteins. The protein sequences of fourteen nicotinic acetylcholine receptors nAChrα1-10 and nAChrβ1-4 were extracted from AaegL5, AEBAN1 and AEBAN2, and aligned using MUSCLE. The phylogenetic tree was constructed using maximum likelihood analysis, with the indicated bootstrap values calculated from 500 replications. The Erwinia ligand-gated ion channel (ELIC) P0C7B7.1 from *Dickeya chrysanthemi* served as the outgroup.

## DISCUSSION

A high-resolution genome sequence assembly of *Aedes aegypti* laboratory strain was reported recently [11]. However, genome sequence information of the wild *Aedes aegypti* population, specially from dengue-endemic region, was not available before. In this study, we report the high-quality consensus genome of *Aedes aegypti* field isolates from Bengaluru, India. Comparative genomic analyses reveal unique genetic differences in the field isolate samples collected from dengue-endemic areas. Based on stringent filtering criteria and using the consensus of three different variant calling tools, we are confident that the genomes of AEBAN1 and AEBAN2 are representative of the variance of natural populations in India. GO analyses showed enrichment of variants in genes encoding membrane proteins and those involved in drug binding and transporter activity. Importantly, our analyses reveal/we find mutations in genes implicated in pesticide resistance. Overall, our observations indicate that, for a given endemic region, comparative phenotypic analyses combined with targeted genomics of a larger *Aedes* sample size will be necessary to determine the optimal vector control strategy for implementation.

## CONCLUSIONS

Our study, for the first time, has generated two high-quality genomes of *A. aegypti* field isolates from southern India, a region endemic to dengue infection. Preliminary comparative genomic analyses highlighted the existence of variations between the laboratory reared *A. aegypti* reference strain LVP_AGWG [11] and wild *A. aegypti* populations in southern India. We believe that this is a useful genomic resource for the mosquito vector genomics community. Future in-depth analyses of additional genomes of field isolates will not only provide insights into the biology of this invasive insect vector but also accelerate research into *Aedes* vector competence, which will help devise rational, effective and sustainable vector transmission control strategies to combat arbovirus infections.

## ACKNOWLEDGEMENTS

The authors thank Lakshmi KJ for help with preparing the genomic libraries. This work was supported by funding from Department of IT, BT & ST, Government of Karnataka, to IBAB and Ramalingaswami re-entry fellowship to SG from the Department of Biotechnology, Government of India. The funding bodies have no role in the design of the study and collection, analysis, and interpretation of data and in writing the manuscript.

