## Supplemental Table S1, Supplemental Figure 1 for "Whole Genome Sequences of *Aedes aegypti* (Linn.) Field Isolates from Southern India"

### Supplementary Material for Bernard *et al.*, 2020

**Supplementary Table S1:** Mapping summary of AEBAN1 and AEBAN2 variants (SNVs and indels) to different genomic regions of the three *A. aegypti* chromosomes

| Sample | Region* | Count | Percent |
|---|---|---|---|
| AEBAN1 | 5' UTR | 114,931 | 0.979% |
| | Exon | 370,253 | 3.155% |
| | Intron | 8,678,342 | 73.95% |
| | 3' UTR | 143,374 | 1.222% |
| | Intergenic | 1,405,422 | 11.976% |
| AEBAN2 | 5' UTR | 37,940 | 0.995% |
| | Exon | 134,201 | 3.518% |
| | Intron | 2,800,268 | 73.414% |
| | 3' UTR | 51,916 | 1.361% |
| | Intergenic | 438,748 | 11.503% |

* The total number of variants for the 3 chromosomes are 3801978 for AEBAN1 and 1210496 for AEBAN2. For several variants, SnpEff assigned more than one effect resulting in 11,735,391 effects for AEBAN1 and 3,814,348 effects for AEBAN2.

**Supplementary Table S2**: Gene Ontology Analysis of variants in the 5'UTR, Exonic and 3'UTR regions of the three chromosomes of *(A)* AEBAN1 and *(B)* AEBAN2 – ***provided as an excel sheet***

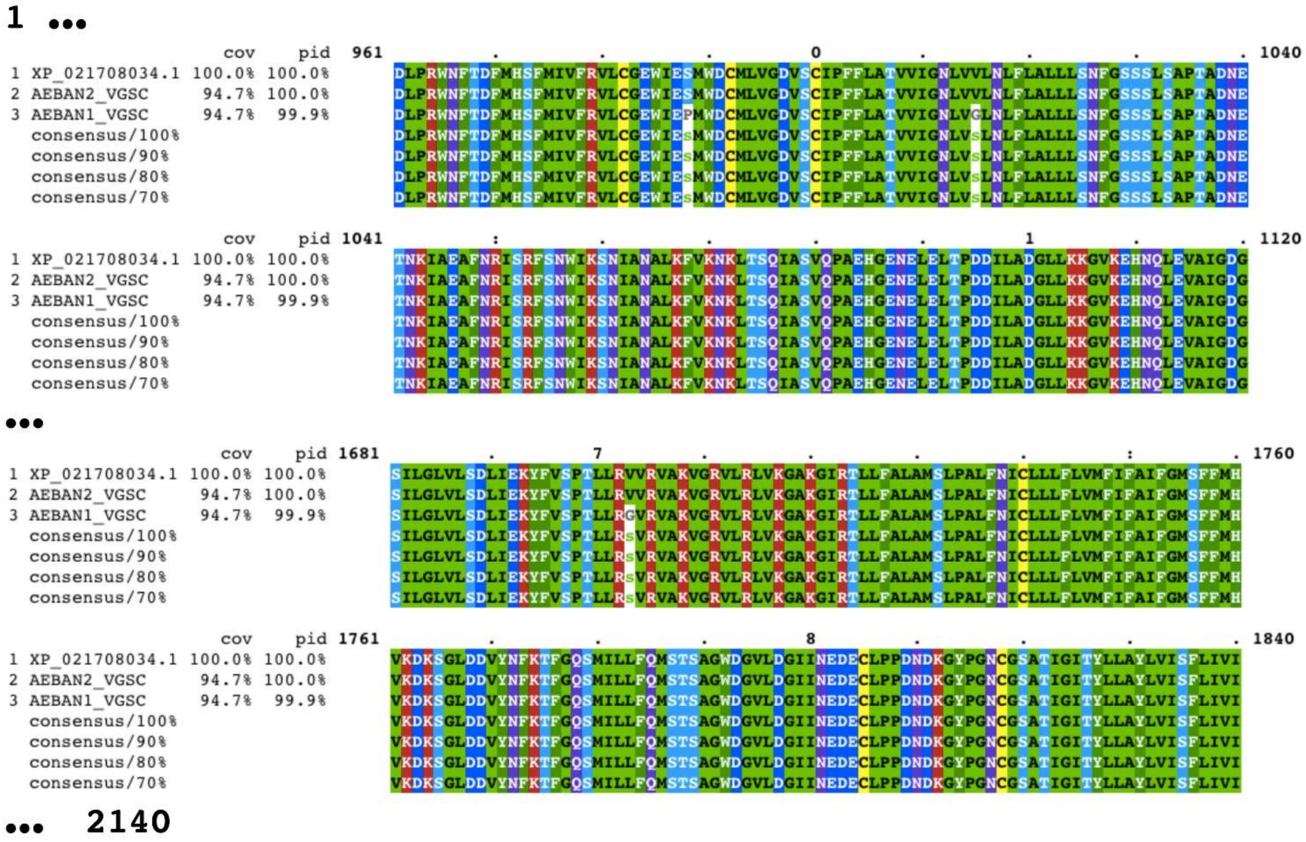

**Supplementary Figure 1: Multiple Sequence Alignment of the voltage gated sodium channel protein (VGSC) from AaegL5, AEBAN1 and AEBAN2.** Protein sequences of the voltage gated sodium channel protein or VGSC from AaegL5, AEBAN1 and AEBAN2 were extracted and aligned using MUSCLE and the alignment visualized using MView. Only the regions that contained amino acid changes in AEBAN1 VGSC are shown here.
